# Ingested insecticide to control *Aedes aegypti*: Developing a novel dried attractive toxic sugar bait device for intra-domiciliary control

**DOI:** 10.1101/794065

**Authors:** Rachel Sippy, Galo E. Rivera, Valeria Sanchez, Froilán Heras Heras, Bianca Morejón, Efraín Beltrán Ayala, Robert S. Hikida, María A. López-Latorre, Alex Aguirre, Anna M. Stewart-Ibarra, David A. Larsen, Marco Neira

**Affiliations:** Institute for Global Health & Translational Science, SUNY-Upstate Medical University, Syracuse, NY, USA; Department of Geography, University of Florida, Gainesville, FL, USA; Center for Research on Health in Latin America, Pontificia Universidad Católica del Ecuador, Quito, Ecuador; Unidad Académica de Ciencias Químicas y de la Salud, Universidad Técnica de Machala, Machala, Ecuador; Department of Medicine and Department of Public Health and Preventative Medicine, SUNY-Upstate Medical University, Syracuse, NY, USA; Department of Public Health, Syracuse University, Syracuse, NY, USA

**Keywords:** *Aedes aegypti*, vector control, toxic sugar bait, attractive bait, semi-field, dengue, arbovirus, ATSB

## Abstract

**Background:** Illnesses transmitted by *Aedes aegypti* (Linnaeus, 1762) such as dengue, chikungunya and Zika comprise a considerable global burden; mosquito control is the primary public health tool to reduce disease transmission. Current interventions are inadequate and insecticide resistance threatens the effectiveness of these options. Dried attractive bait stations (DABS) are a novel mechanism to deliver insecticide to *Ae. aegypti*. The DABS are a high-contrast 28 inch^2^ surface coated with dried sugar-boric acid solution. *Ae. aegypti* are attracted to DABS by visual cues only, and the dried sugar solution elicits an ingestion response from *Ae. aegypti* landing on the surface. The study presents the development of the DABS and tests of their impact on *Ae. aegypti* mortality in the laboratory and a series of semi-field trials.

**Methods:** We conducted multiple series of laboratory and semi-field trials to assess the survivability of *Ae. aegypti* mosquitoes exposed to the DABS. For laboratory experiments we assessed the lethality, the killing mechanism, and the shelf life of the device through controlled experiments. In the semi-field trials, we released laboratory-reared female *Ae. aegypti* into experimental houses typical of peri-urban tropical communities in South America in three trial series with six replicates each. Laboratory experiments were conducted in Quito, Ecuador, and semi-field experiments were conducted in Machala, Ecuador – an area with abundant wild populations of *Ae. aegypti* and endemic arboviral transmission.

**Results:** In the laboratory, complete lethality was observed after 48 hours regardless of physiological status of the mosquito. The killing mechanism was determined to be through ingestion, as the boric acid disrupted the gut of the mosquito. In experimental houses, total mosquito mortality was greater in the treatment house for all series of experiments (p<0.0001).

**Conclusions:** The DABS devices were effective at killing female *Ae. aegypti* under a variety of laboratory and semi-field conditions. DABS are a promising intervention for interdomiciliary control of *Ae. aegypti* and arboviral disease prevention.

## Background

Arboviral illnesses, including dengue, chikungunya, yellow fever, and Zika, are major contributors to morbidity and mortality in the tropics and subtropics. The burden is particularly apparent in Central and South America: between 2010—2018, the estimated annual number of dengue cases in the region ranged from 500,000 to 2,400,000 [1], and since 2013 PAHO has estimated that there have been more than 2.5 million suspected and confirmed cases of chikungunya and 800,000 cases of Zika. The viruses causing these diseases are spread mainly by the mosquitoes *Aedes aegypti* (Linnaeus, 1762; *Ae. aegypti*) and *Aedes albopictus* (Skuse, 1894; *Ae. albopictus*), with *Ae. aegypti* serving as the principal vector in many South American countries, including Ecuador [2]. Due to the lack of commercially-available vaccines for most human arboviral diseases, prevention efforts focus on vector surveillance and control methods [3].

Vector control relies heavily on contact-based insecticides, which are available in four main classes: organophosphates, pyrethroids, carbamates, and organochlorines. Indoor residual spraying is a common approach to vector control, for which twelve insecticides are available and approved for human use [4]. This small number of approved insecticides constitutes an impediment for the implementation of effective vector control strategies (such as pesticide rotation cycles) aimed at decreasing the development of resistance to any single insecticide [5]. As a result, pesticide resistance has become a major limitation for current vector control strategies, and is widespread in South American countries [6–8]. Our current reliance on a few chemical molecules to control *Ae. aegypti* is an increasingly flawed strategy, as evidenced by the proliferation of this disease vector across the globe and increasing arbovirus epidemics [9].

In contrast to the contact-based insecticide approach of the public health sector, the agricultural industry has focused on ingested insecticides for pest control. The use of ingested insecticides could be applied in disease control programs and interventions if disease vectors are successfully led to ingest the insecticide. One solution, attractive toxic sugar baits (ATSB), exploits the nectar-feeding behavior of mosquitoes [10,11] to deliver the insecticide. An ATSB uses a mixture of a lethal agent with sugar water and an additional attractant [12]. ATSBs have been tested for *Anopheles* spp. [13–17], *Culex* spp. [15,16,18,19], *Ae. albopictus* [20–23], and other vector or nuisance species [16] with a variety of attractants, baits, active ingredients, designs, and placement strategies. Although laboratory bioassays demonstrate that ATSBs are toxic to *Ae. aegypti* [16,24,25], semi-field and field evaluations have had poor results in reducing *Ae. aegypti* populations [26,27], indicating that ATSB devices must be carefully designed and tested for each target species [12].

Compared to other mosquito species, *Ae. aegypti* appear to have a lower propensity for sugar-feeding, preferring human blood meals instead [11]. Despite this, *Ae. aegypti* females will readily feed on sugar in the laboratory, and often feed on plant sugars in the wild [28–31]. However, traditional attractive sugar bait strategies that rely only on fruit volatiles as an attractant are likely insufficient to “lure” highly anthropophilic female *Ae. aegypti* in the natural environment.

Herein we present the development of dried attractive bait stations (DABS) (Figure 1), and show results from laboratory and semi-field experiments. In the laboratory we first identified the lethality of DABS (Series 1.1), aimed to identify the killing mechanism of the DABS (Series 1.2), assessed how the physiological status altered the effectiveness of DABS (Series 1.3), and assessed the shelf life of the DABS (Series 1.4). In the semi-field trials, we sought to determine the timing of mosquito mortality (Series 2.1), assess the relationship between DABS exposure time and mosquito mortality (Series 2.2), and to demonstrate these effects in the presence of competing attractants (Series 2.3).

**Figure 1.**
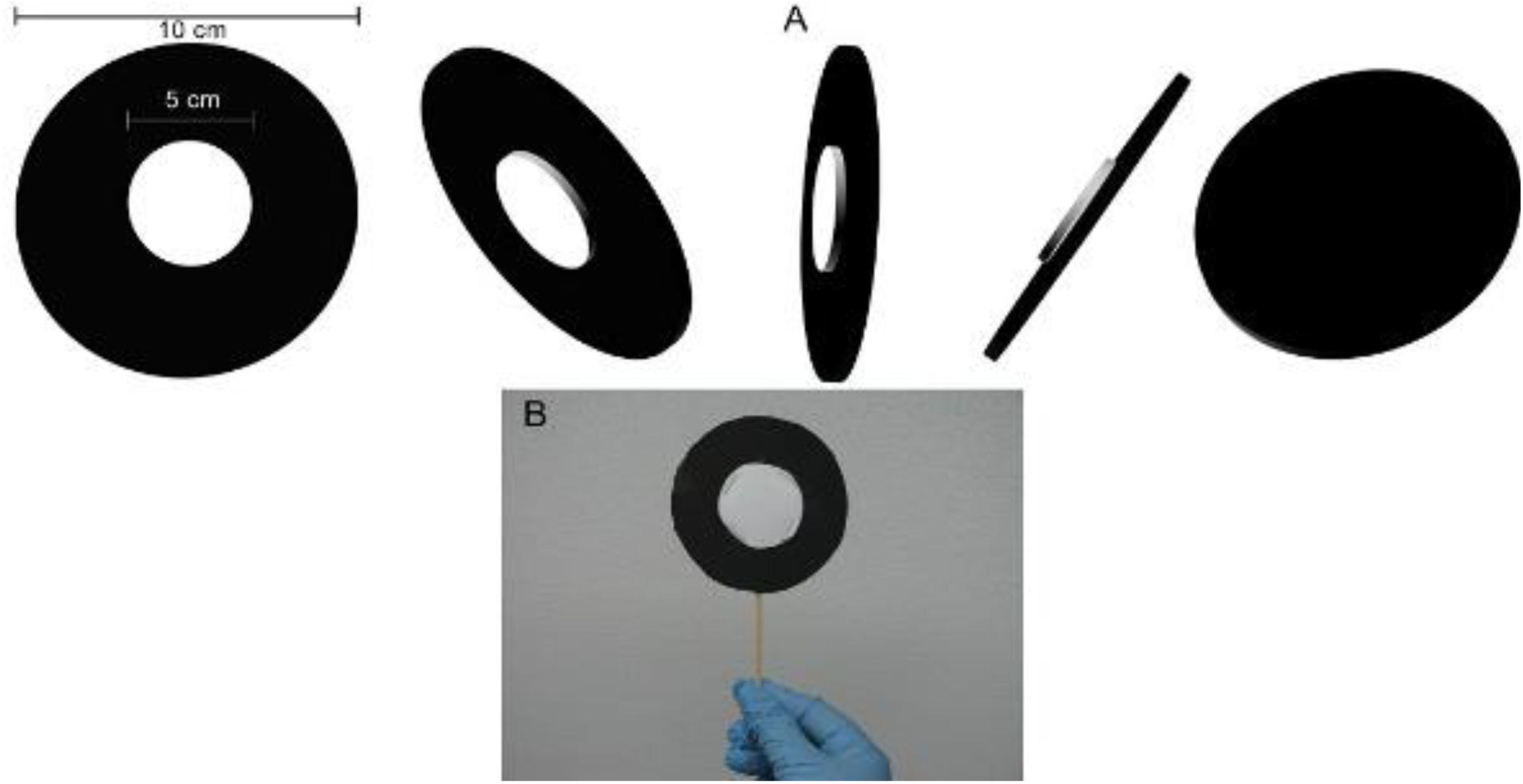
Attractive toxic sugar bait devices. A) 3D model of the DABS devices. Diameters for each of the foam sheet circles are shown.B) Photography of a DABS device used in the study.

## Results

### Laboratory experiments

#### Series 1.1: Effects of DABS exposure on mosquito survival

We measured survival in mosquitoes exposed to DABS and compared to mosquitoes exposed to control DABS in 20×20×20cm cages during in four independent replicates. An average of 13.5 (N=4, std = 3.88, SE= 1.94) out of 30 mosquitoes exposed to DABS survived at the first 24 hours post-exposure. All mosquitoes had died by the end of the experiments at 48h post-exposure (Figure 2). In contrast, an average 29.75 (N=4, std = 0.50, SE=0.25) out of 30 mosquitoes exposed to control devices survived after the first 24h. After the second 24h period, by the end of the experiment an average of 29. 25 (N=4, std= 0.96, SE=0.48) had survived. Differences between toxic and control treatments were highly significant at 24h (df=7, t=8.32, p<0.001) and 48h (df=7, t=61.1, p<0.001) post-exposure.

**Figure 2.**
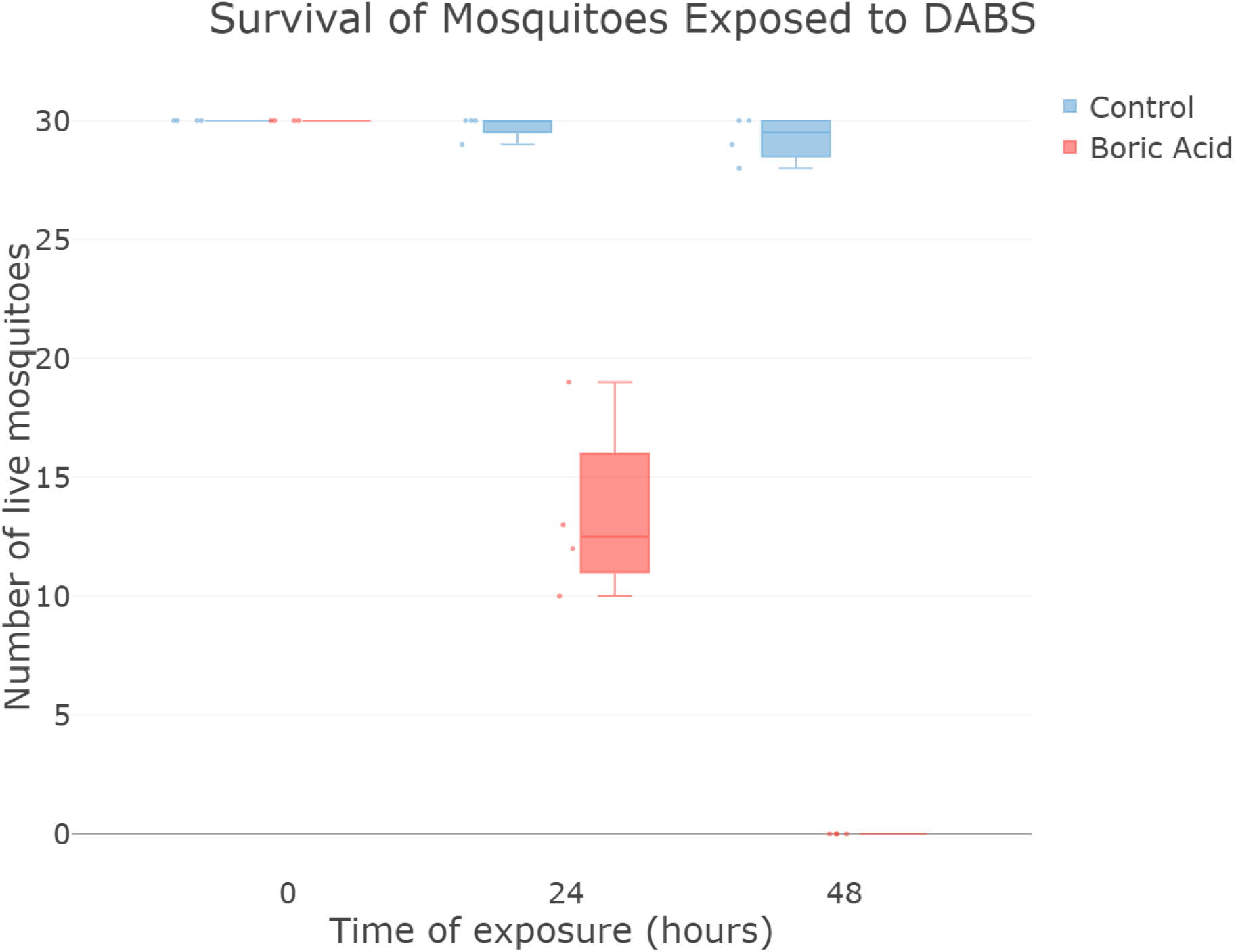
Survival assessment of mosquitoes exposed to the device. All mosquitoes (n=30) exposed to toxic devices die after 48 hours of exposure. When presented with non-toxic device almost all survive. Box plot indicating median 25% and 75% quartiles. Error bars indicate max and min values. Each dot indicates an independent count of mosquitoes (y axis) exposed to control devices (blue) or toxic devices (red) at different time points (x axis).

#### Series 1.2: Characterization of the biological mode of action of the device

We disrupted the feeding parts of mosquitoes and examined survival in those exposed to DABS compared to those exposed to control DABS. After 48 hours, all mosquitoes that could still feed (*i.e*. mosquitoes with an intact proboscis), died when exposed to the toxic device, and out of 20 19.33 (N=3, std= 0.58, SE=0.29) survived when exposed to the non-toxic control device. On average 12.33 mosquitoes that could not feed (those with ablated proboscis), survived regardless of inclusion of the device’s toxic condition (N=3, std=1.52, SE=0.87) or control non-toxic condition (N=3, std=2.86, SE=1.65). Significant differences were found between the four treatments (df=11, F=70.55, p<0.001). Post-hoc pairwise comparison determined that (a) the mortality of ablated mosquitoes exposed to toxic devices was not significantly different to the mortality of ablated mosquitoes exposed to control devices, and (b) the mortality of ablated mosquitoes was significantly different from the mortality of whole mosquitoes exposed to toxic devices and whole mosquitoes exposed to control devices (Figure 3).

**Figure 3.**
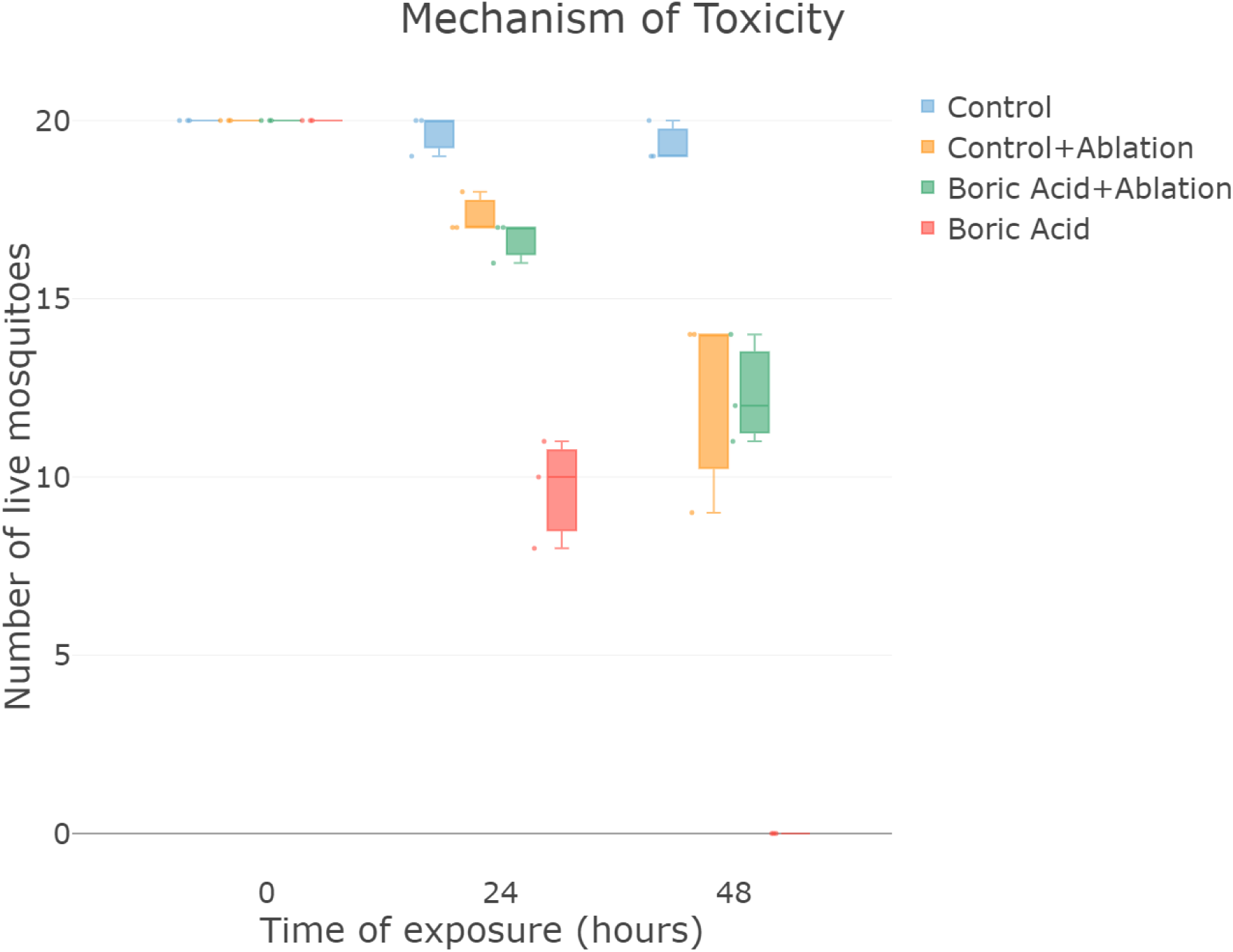
Uptake mechanism of the toxic component. Toxic effect is dependent on the ability of mosquitoes to ingest the toxic component. When mosquitoes are able to ingest the toxic component all mosquitoes (n =20) die after 48 hours (red). Mosquitoes with ablated mouthparts die equally regardless of the toxic or non-toxic condition of the device (green and yellow). Almost all mosquitoes with no ablation or toxic devices survive. Box plot indicating median 25% and 75% quartiles. Error bars indicate max and min values. Each dot indicates an independent count of mosquitoes (y axis) exposed to control devices (blue), control devices + mosquito mouthparts ablation (yellow), toxic devices + mosquito mouthparts ablation, or toxic devices (red) at different time points (x axis).

Mosquitoes that had ingested toxic sugar solution presented histological abnormalities in the posterior midgut. Electron micrographs revealed disruptions in the continuity of the gut epithelium (Figures 4A, 4C). In addition, we found abnormal adipocytes that we hypothesize are undergoing a process of necrosis (Figures 4E, 4F), and an increase in both the size and number of basal infolds in the gut epithelial cells. We hypothesize that boric acid ingestion is the cause of these pathological changes, which contributed to the mortality observed in specimens exposed to toxic devices. Microscopic images of individuals exposed to control devices presented none of these pathologies on the posterior midgut (Figures 4B, 4D).

**Figure 4.**
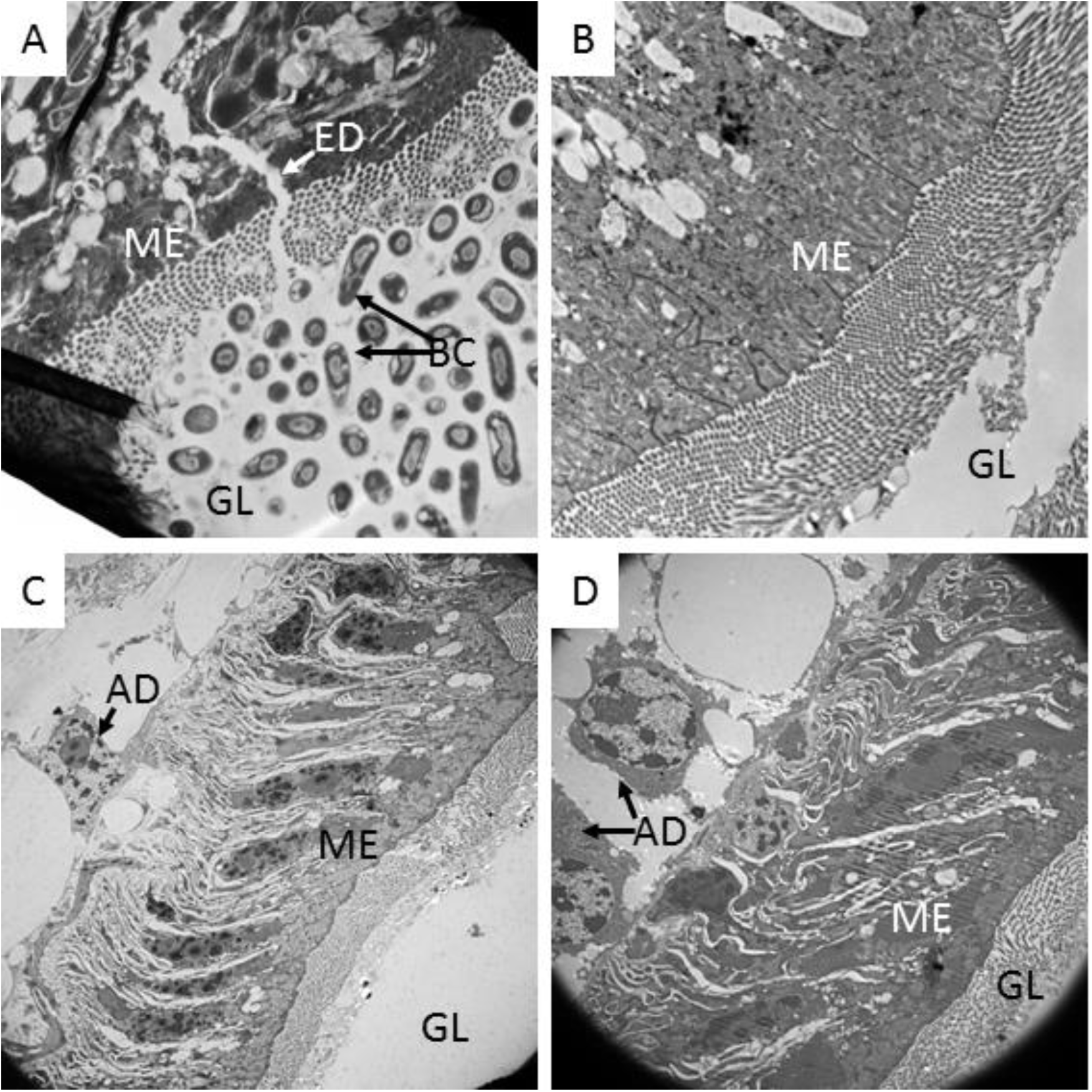
Histopathological effects on the midgut. Longitudinal sections of Ae. aegypti posterior midgut. Panels A, C, D: Mosquitoes exposed to toxic devices. Panel B: Mosquito exposed to non-toxic device. Specimens exposed to toxic devices showed disruptions in the gut integrity (ED, panel A). Because of the even distribution of adjacent bacterial cells in the gut lumen, this disruption is unlikely to be the result of sample processing for electron microscopy. Abbreviations: AD: Adipocite; BC: Bacterial cells in gut lumen; ED: Epithelial disruption; GL: Gut lumen; ME: Midgut epithelium. Magnifications: A: 15,000×; B: 10,000×; C: 3,000×; D: 5,000×.

#### Series 1.3: assessment of mosquito physiological status on DABS effectiveness

We measured survival in mosquitoes at blood fed and parous life stages in mosquitoes exposed to DABS and mosquitoes exposed to control DABS. Mosquitoes with both physiological statuses evaluated (blood fed and parous) presented lower survival when exposed to toxic devices than when exposed to non-toxic control devices. In toxic conditions, an average of 19.33 (N= 3, std= 1.73, SE=0.99) blood fed females’ out of 30 had survived at 48h. 2.67 (N= 3, std= 3.05, SE=1.76) blood fed mosquitoes had survived at 72h, by the end of the experiment. 27 (N= 3, std= 1.73, SE=0.99) out of 30 blood fed mosquitoes had survived to non-toxic DABS at the end of 72 hours (Figure 5). Differences between control and toxic treatment survival were significant at 48h (df=5, t=5.75, p<0.01) and 72h (df=5, t=12, p<0.001) post-exposure.

**Figure 5.**
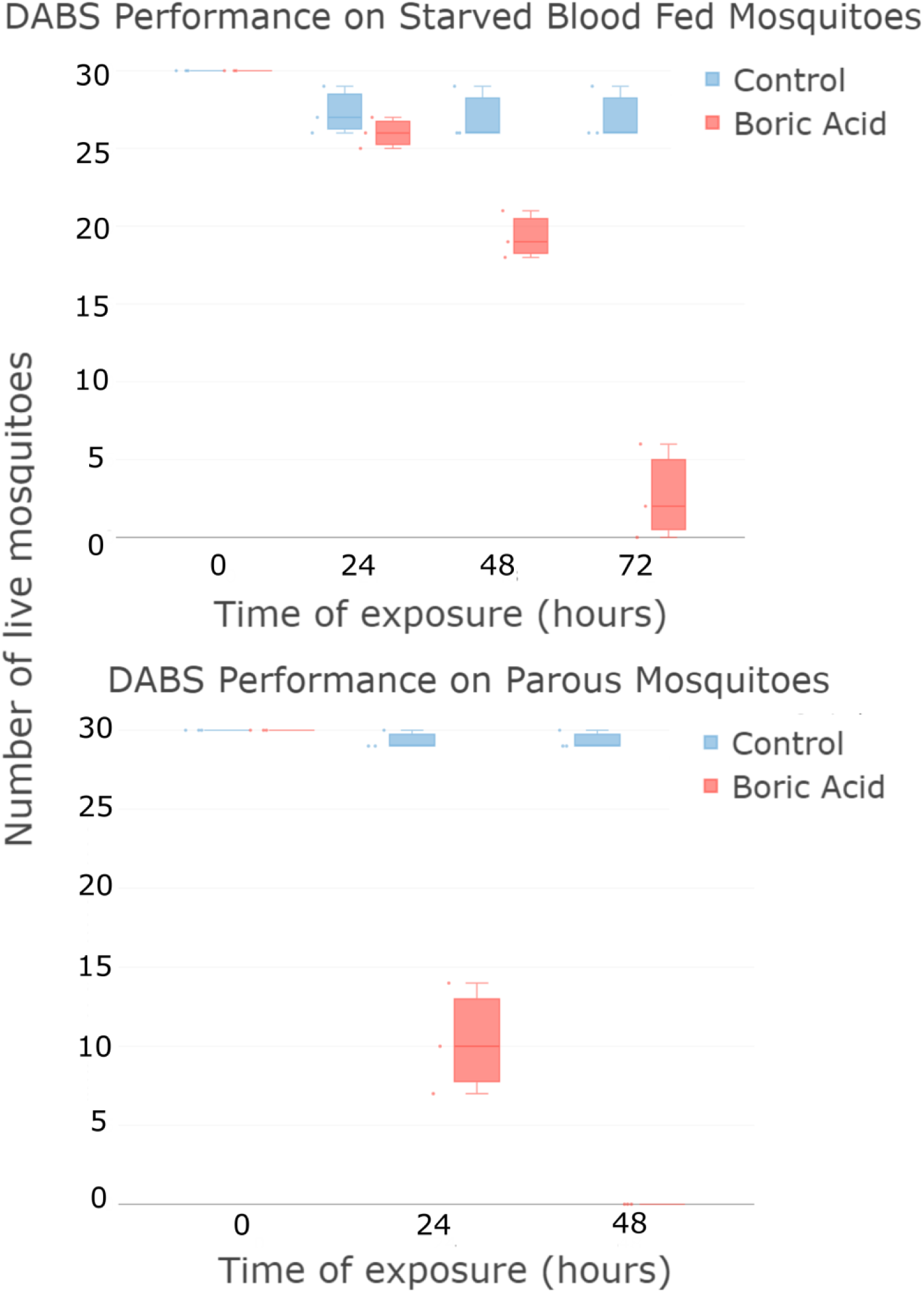
Effects of the physiological status of the mosquitos on the performance of the device. DABS are efficient at killing mosquitoes through different physiological stages. (A) A significant majority of blood fed mosquitoes exposed to toxic devices die after 72 hours of exposure. When presented with non-toxic device almost all survive. (B) All parous mosquitoes exposed to toxic devices die after 48 hours of exposure. When presented with non-toxic device almost all survive. Box plot indicating median 25% and 75% quartiles. Error bars indicate max and min values. Each dot indicates an independent count of mosquitoes (y axis) exposed to control devices (blue) or toxic devices (red) at different time points (x axis).

10.33 (N= 3, std= 3.51, SE=2.02) parous female mosquitoes at the first 24 hours and a 0 at 48 hours of exposure had survived to the toxic DABS devices (Figure 5B). In the non-toxic control group 29.33 (N= 3, std= 0.58,) at 48 hours of exposure. Differences between control and toxic treatment survival curves were significant at 24h (df=5, t=9.25, p<0.001) and 48h (df=5, t=87.99, p<0.001) post-exposure.

#### Series 1.4: Assessment of shelf-life of the DABS device

We tested the shelf life of DABS by measuring survival of mosquitoes exposed to DABS compared to those not exposed to DABS, using DABS that had been constructed at various time points in the past. 30 out of 30 mosquitoes exposed to toxic devices stored for 38 days had survived at 24 hours; 28.67 (N= 3, std= 0.58, SE=0.33) mosquitoes exposed to control conditions survived at 48 hours (Fig. 6A). Differences in survival between conditions were highly significant at 48h post-exposure (df=5, t=86, p < 0.001).

**Figure 6.**
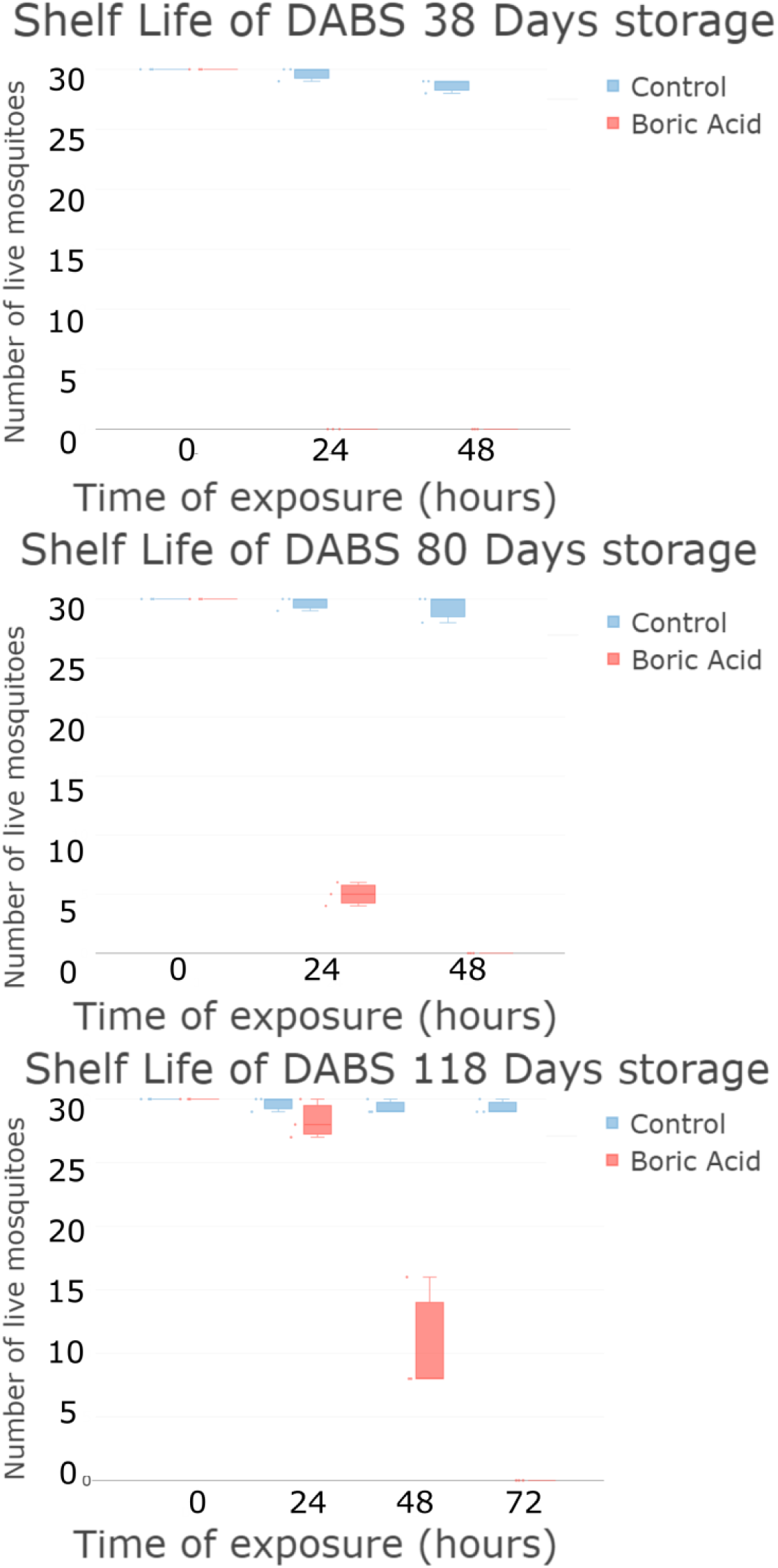
Shelf life of the device. DABS are still effective after (A) 38 days, (B) 80 days and (C) 118 days of storage in insectary conditions. Box plot indicating median 25% and 75% quartiles. Error bars indicate max and min values. Each dot indicates an independent count of mosquitoes (y axis) exposed to control devices (blue) or toxic devices (red) at different time points (x axis).

5 (N= 3, std= 1.0, SE=0.58) mosquitoes exposed to toxic devices stored for 80 had survived at the first 24 hour period and 0 after 48 hours; 29.33 (N=3, std=1.16, SE= 0.67) mosquitoes exposed to control conditions survived at 48 hours (Fig. 6B). Differences in survival between conditions were highly significant at 48h post-exposure (df=5, t=44, p < 0.001).

On average 28.33 (N=3, std= 0.58, SE=0.33), 10.66 (N=3, std=4.62, SE=2.67), and 0 mosquitoes exposed to toxic devices stored for 118 had survived at 24h, 48h, and 72h, respectively (Fig. 6C). Differences in survival between conditions were highly significant at 48h (df=5, t=6.95, p<0.01) and 72h (df=5, t=87.99, p < 0.001) post-exposure.

### Semi-field experiments

We assessed the attractiveness of DABS by measuring mortality in mosquitoes exposed to DABS compared to mosquitoes not exposed to DABS in experimental houses. When exposed to DABS in semi-field trials (Series 2.1, Figure 7), mosquito mortality was 0.0—6.0% (mean: 2.0%, standard error: 0.9%) in the control and 17.0—57.1% (mean: 36.7%, standard error: 5.3%) in the treatment house after 24 hours (df=5, t=-7.0, p<0.001). At 48 hours, mortality was 0.0—18.0% (mean: 5.4%, standard error: 2.4%) in the control and 22.0—51.1% (mean: 38.9%, standard error: 3.9%) in the treatment house (df=5, t=-5.36, p<0.01). At 72 hours, mortality was 0.0—4.1% (mean: 0.7%, standard error: 0.6%) in the control and 0.0—4.0% (mean:1.4%, standard error: 0.6%) in the treatment house (df=5, t=-0.80, p>0.05). The cumulative mortality of the control was 4.1—18.0% (mean: 8.2%, standard error: 1.9%) and 54.0—98.0% (mean: 76.9%, standard error: 6.2%) in the treatment house (df=5, t=-8.37, p<0.001). Most mosquito mortality was observed within the first 48 hours of the experiment, with no difference in mosquito mortality after this time period.

**Figure 7:**
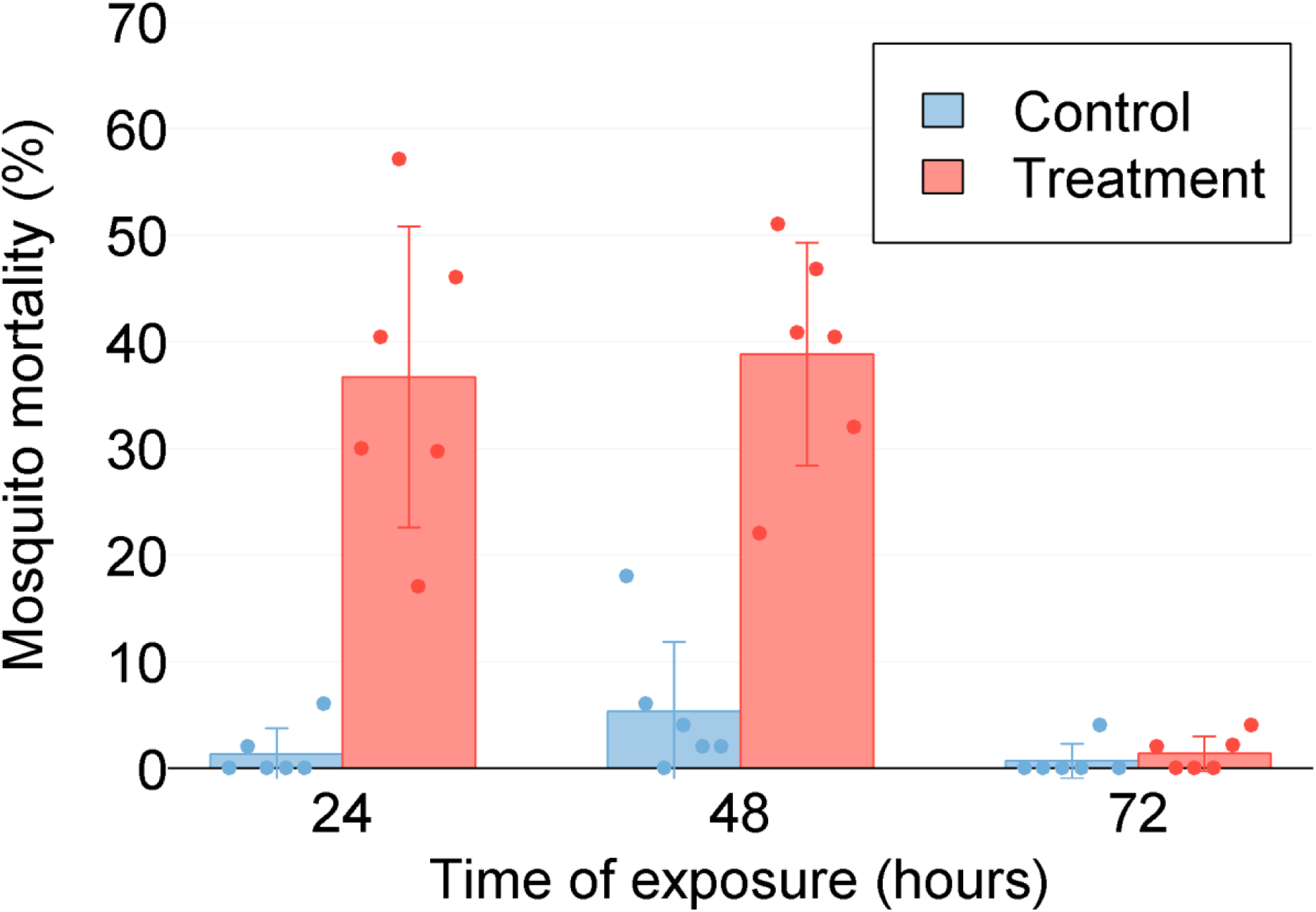
Mortality of mosquitoes when exposed to DABS over time (Series 2.1). Mosquitos were exposed to DABS for 24 hours and observed afterwards; mosquito mortality was calculated immediately after the exposure period, and during laboratory observation at 48 hours and 72 hours. Mean control and experimental house mortalities are shown as bars, and standard deviation as error lines. Points indicating the mortality from each individual replicate are overlaid on each experimental condition.

When exposed to DABS for 48 hours (Series 2.2, Figure 8), mosquito mortality was 2.0— 22.9% (mean: 11.7%, standard error: 2.8%) in the control and 77.3—100.0% (mean: 91.5%, standard error: 3.8%) in the treatment house (df=5, t=-17.0, p<0.001), indicating high mortality from 48 hours of exposure to DABS in the treatment houses.

**Figure 8:**
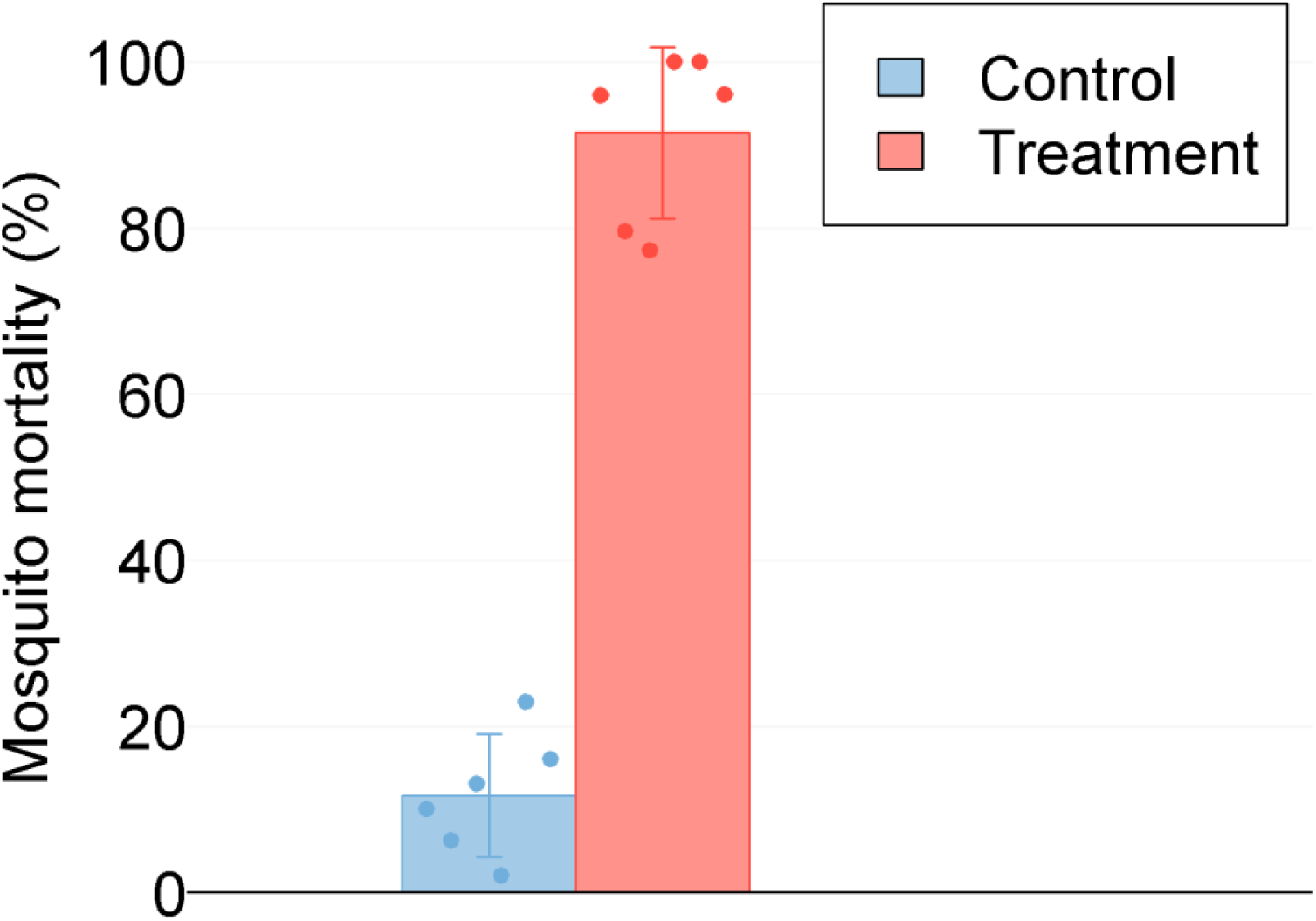
Mortality of mosquitoes when exposed to DABS for 48 hours (Series 2.2). Mosquitos were exposed to DABS for 48 hours; mosquito mortality was calculated immediately after the exposure period. Mean control and experimental house mortalities are shown as bars, and standard deviation as error lines. Points indicating the mortality from each individual replicate are overlaid on each experimental condition.

When alternative attractants were included in the experimental houses (Series 2.3, Figure 9), mosquito mortality was 2.0—32.7% (mean: 14.1%, standard error: 4.1%) in the control and 68.0— 100.0% (mean: 89.6%, standard error: 4.5%) in the treatment house (df=5, t=-12.90, p<0.001), indicating that DABS results in high mortality even in the presence of a competing attractant. When comparing the results of 24 hours (Series 2.1) to 48 hours of exposure (Series 2.2), 48 hours of exposure results in higher mortality at 48 hours (df=10, t=-8.78, p<0.001) in the treatment group (Supplemental Table 1), with no difference in the control groups (df=10, t=-1.55, p>0.05).

**Figure 9:**
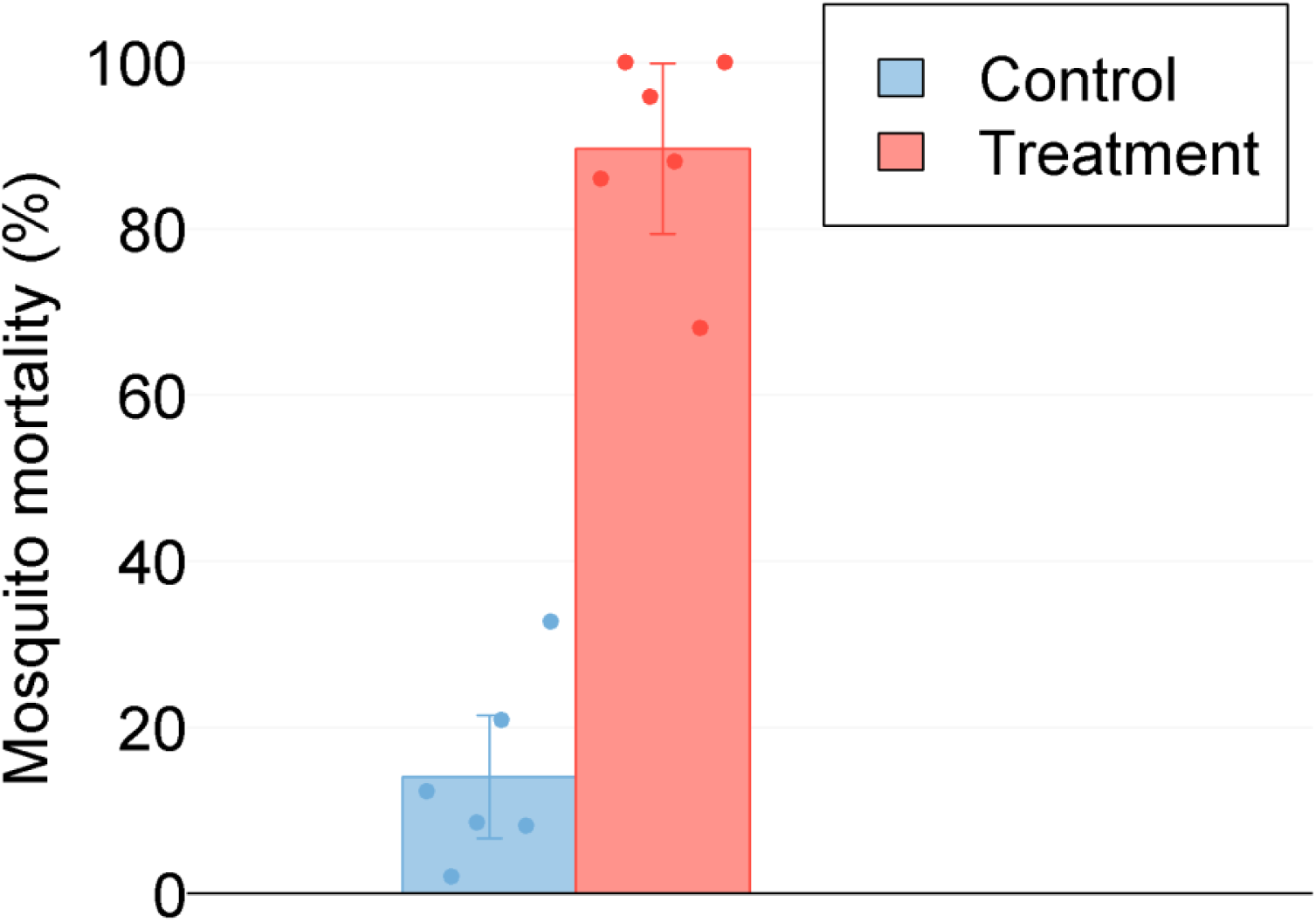
Mortality of mosquitoes when exposed to DABS for 48 hours in the presence of alternative sugar source (Series 2.3). Mosquitos were exposed to DABS for 48 hours in the presence of a competing attractant; mosquito mortality was calculated immediately after the exposure period. Mean control and experimental house mortalities are shown as bars, and standard deviation as error lines. Points indicating the mortality from each individual replicate are overlaid on each experimental condition and time point.

When comparing 48 hours of exposure to DABS only (Series 2) and 48 hours of exposure to DABS in the presence of a competing attractant (Series 2.3), there is no effect of a competing attractant on the effect of DABS on mosquito mortality (df=10, t=0.28, p>0.05) in the treatment group (Table 1). High mortality from 48 hours of DABS exposure was observed despite the presence of a competing attractant.

**Table 1:** Comparison of Results Across Semi-field Experimental Series. Mean and (standard error) of 48-hour mosquito mortality from 6-replicate series are given. Mean mortality was compared across trial series using a t-test.

| Table 1: Comparison of Results Across Semi-field Experimental Series |  |  |  |
| --- | --- | --- | --- |
| Mean and (standard error) of 48-hour mosquito mortality from 6-replicate series are given. Mean mortality was compared across trial series using a t-test. |  |  |  |
| Experimental Conditions |  | 48-hour Mortality | p-value |
| Control | 24-hour DABS exposure (Series 1) | 5.5% (2.4) | 0.151 |
|  | 48-hour DABS exposure (Series 2) | 11.7% (2.8) |  |
| Treatment | 24-hour DABS exposure (Series 1) | 38.9% (3.9) | <0.001 |
|  | 48-hour DABS exposure (Series 2) | 91.5% (3.8) |  |
| Control | 48-hour DABS exposure (Series 2) | 11.7% (2.8) | 0.671 |
|  | 48-hour DABS exposure with competing attractant (Series 3) | 14.1% (4.1) |  |
| Treatment | 48-hour DABS exposure (Series 2) | 91.5% (3.8) | 0.784 |
|  | 48-hour DABS exposure with competing attractant (Series 3) | 89.6% (4.5) |  |

## Discussion

These experiments demonstrate that DABS can strongly impact the mortality of female *Ae. aegypti* under laboratory and semi-field conditions. In these settings, we show that mortality occurs within the first 48 hours of exposure to our devices. In addition, DABS attract and kill *Ae. aegypti* even in the presence of an alternative sugar source. To the best of our knowledge, this device is the only known “dry” ATSB. The simple and economic design lends itself to in-home use in resource limited settings where *Ae. aegypti* target human hosts and transmit dangerous arboviruses.

Our assessment of the biological action of the devices provides an insight into the mechanism by which low concentrations of boric acid affect *Ae. aegypti*. We determined that boric acid enters the insect’s body by ingestion, further supporting the notion that this inorganic pesticide acts as a stomach poison, as previously suggested [32,33]. Based on our electron microscopy analysis, we hypothesize that the ingestion of boric acid disrupts the integrity of the gut epithelium.

Considering that the proposed mechanism by which boric acid exerts its toxic effect (gut disruption) is markedly different from the neurotoxic mechanism by which most traditional pesticides cause mortality, we propose that our devices have the potential to act as efficient complementary tools to combat the spread of resistance to traditional pesticides. By combining the use of DABS with traditional pesticides in the same areas, it would be possible to target two different and crucial systems (namely, the nervous and digestive systems) in the insect’s body simultaneously, thereby reducing the mosquito’s probability of survival and decreasing the probability of the development of insecticide resistance.

We observed significant mortality of blood fed female *Ae. aegypti* exposed to the DABS device, albeit at lower rate than starved females. Interestingly, the largest drop in survival probability in blood fed females is observed between 48h and 72h post-exposure to the device (Fig 5), suggesting that after 48h, females have already used imbibed blood for the development of eggs and are keen to search for further meals. Based on this evidence, it is plausible to suggest that if employed in the field, DABS devices may be efficient in killing female mosquitoes of various physiological states, including females that have already ingested blood – a particularly important group for disease transmission.

Novel vector control methods have the potential to serve as critical tools in the public health effort to control persistent and emerging vector borne diseases. Various designs of ATSBs have had promising field trials for potential control of *Aedes albopictus* Skuse, 1894, *Anopheles* spp. and *Culex* spp. [15–17,20,21,24]. Previous research shows that several formulations of ATSBs can achieve *Ae. aegypti* mortalities above 80% in laboratory settings [16,25], but results from ATSBs in semi-field or field settings have been mixed. Early field trials did not show a positive effect of ATSBs on *Ae. aegypti* [26,27], however a recent field trial in Bamako, Mali showed promising success [31]. The principle barrier to field trial success appears to be the ability to attract *Ae. aegypti* to ATSBs and mixed results have been achieved when using floral-based attractants.

We hypothesize that our device attracts *Ae. aegypti* with strong visual cues (as opposed to a chemical) as an attractant. *Ae. aegypti* are container breeders [34,35], that utilized tree holes in their natural forested habitat before adapting to life in human civilization. The DABS device has a high-contrast (black and white) 28-inch^2^ surface to simulate a refuge for *Ae. aegypti* [36]. High contrast coloring has similarly been integrated into prior trap designs and has been shown to improve capture rates of *Ae. aegypti* [37]. We believe the high-contrast coloring of DABS draws *Ae. aegypti* to land on the device.

These experiments have demonstrated the effectiveness of DABS on *Ae. aegypti* in laboratory and semi-field experimental conditions. Our approach differs from most ATSB approaches in two important ways: firstly, we use a device with a dried sugar solution to elicit an ingestion response while other ATSBs typically use liquid sprayed on vegetation [12,15,17,26]. We hypothesize that the device is a key element in the effectiveness of DABS. Similar to other diptera [38], *Ae. aegypti* are able to evaluate surfaces with their feet, and the “taste” of a landing surface can either lead the mosquito to feed and ingest, or reject, the surface [39]. Additionally, the device provides two operational advantages over spraying liquid solutions: 1) liquid solutions are more difficult to manufacture, ship, and distribute than devices, and 2) the device can be smaller and more easily deployed. Secondly, we use a visual rather than chemical attractant to lure *Ae. aegypti* to the device. Chemical attractants add to the cost and decrease the shelf life of any device. Previous research has questioned the ability of sugar solutions alone to attract mosquitoes [26,33], leading to research on chemical attractant additives for ATSBs, but the use of chemical attractants in ATSBs targeting *Ae. aegypti* have been unsuccessful [26,27]. We demonstrate that a simple black-and-white visual attractant is a sufficient motivator for female *Ae. aegypti* to land on the surface of DABS even in the presence of a competing oasis. Taken together, we hypothesize that the visual cues attract the *Ae. aegypti* to land on the device, upon which the presence of the dry sugar on the device’s surface entices the insect to ingest it. When this sugar solution is mixed with boric acid, ingestion results in insect mortality.

We propose that these encouraging results justify larger field trials of DABS in open-air environments. We show that 48 hours of DABS exposure leads to high mosquito mortality when used in the laboratory and in experimental houses reminiscent of peri-urban tropical housing. Furthermore, we have established that the effectiveness of DABS for killing *Ae. aegypti* is maintained even after prolonged storage periods, a characteristic that would facilitate their use in semi-field and field conditions.

Semi-field trials are a crucial step to bring a scalable, marketable product to intra domiciliary field testing. An in-home approach is ideal for control of *Ae. aegypti*, as the vector has an extremely limited flight range, often spending its entire life within a single household [5,35,40]. Other research with ATSBs has shown that end-users of these products prefer to have them placed indoors [14]. The successful design and placement strategy of DABS used in our experiments indicate that the device is ideal for in-home field testing.

### Limitations

These experiments were conducted under laboratory and semi-field conditions, which can only moderately emulate real-world/field conditions. Semi-field experiments were limited to nulliparous females and we cannot be certain how DABS will affect gravid or blood-fed females or males in an open-air environment—though it should be noted that DABS were equally effective in attracting and killing blood-fed and nulliparous females under laboratory conditions. It is also unclear if DABS would impact non-target insect species, such as butterflies or other pollinators, though if DABS are limited to use inside the home, it is unlikely to affect these species. Although DABS performed well in the presence of a competing attractant (100 grams of apples), it is unlikely that the attractant used in our experiments are a realistic substitute for open-air field conditions. An actual home will contain many competing attractants, including human hosts. It is difficult to know if the success of DABS in semi-field conditions will be replicated in occupied homes in the field; the number and placement of DABS may need to be modified. In addition, it is unclear how end users will react to placement of DABS in their homes, although our preliminary examinations (unpublished) suggest residents are receptive of DABS and there is evidence that residents in areas of high *Ae. aegypti* burden are willing to utilize numerous home-based mosquito control products [41].

### Conclusions

With careful design and device placement consideration, we have created a promising vector control device ready for large-scale trials to test its ability to control *Ae. aegypti* in natural conditions. We demonstrated that DABS are capable of attracting and killing female *Ae. aegypti* in experimental houses, and that 48 hours in the presence of DABS leads to high mortality among female *Ae. aegypti*. Importantly, DABS were efficient at killing female mosquitoes of diverse physiological statuses, and can attract and kill female *Ae. aegypti* even in the presence of a competing attractant.

## Materials and methods

### Aim and design

The general aim of our study was to assess the effectiveness of DABS at killing female *Ae. aegypti* in numerous laboratory and semi-field experimental conditions. In the laboratory we first measured a killing effect of DABS (Series 1.1), identified the killing mechanism (Series 1.2), and measured how mosquito physiological status affected DABS lethality (Series 1.3). In the semi-field experiments our specific aims were (a) to determine the timing of mosquito mortality (Series 2.1), (b) to assess the relationship between DABS exposure time and mosquito mortality (Series 2.2), and (c) to demonstrate these effects in the presence of competing attractants (Series 2.3).

In the laboratory we first identified the lethality of DABS (Series 1.1), aimed to identify the killing mechanism of the DABS (Series 1.2), assessed how the physiological status altered the effectiveness of DABS (Series 1.3), and assessed the shelf life of the DABS (Series 1.4). In the semi-field trials, we sought to determine the timing of mosquito mortality (Series 2.1), assess the relationship between DABS exposure time and mosquito mortality (Series 2.2), and to demonstrate these effects in the presence of competing attractants (Series 2.3).

### Study setting

#### Laboratory experiments

Laboratory experiments were conducted at the Center for Research on Health in Latin America (CISeAL, by its Spanish acronym), where they were reared and maintained under standard insectary conditions: 28 ± 1°C temperature, 80 ± 10% relative humidity, and a 12h:12h (L:D) light cycle. Larvae were fed finely ground fish food. When required, mosquitoes were sexed during the pupal stage. Adults were kept in 20×20×20 cm cages. For maintenance, adult mosquitoes were fed 10% sucrose solution *ad libitum*. For blood feeding, female adult mosquitoes were offered access to a restrained female mouse. All mosquitoes were maintained under insectary conditions after adult emergence before they were used for experiments. Mosquitoes referred to as “starved” in this manuscript were deprived of access to sugar or blood (but not water) for 48h prior to their use in experiments.

#### Semi-field trials

Assays were performed in experimental houses meant to emulate typical housing found in areas with active dengue transmission. Photographs of the houses are available in an additional file (see Additional file 1). The houses are constructed of wood and cane and are raised on a 1-meter platform with walkways to improve structural integrity and facilitate window access; one window on each house is equipped with window escape traps with sleeves to monitor escape behavior. The dimensions of the houses are 3.85 meters (m) wide x 4.85 m long x 3 m high. Each house has three windows (0.9 m wide × 0.6 m high) and one door (1.03 m wide × 3 m high). The house frames are made of wood, untreated wooden plank flooring, walls of untreated cane, and a roof of zinc panels. The window traps are 0.45 m long x 0.66 m wide x 0.45 m high. The houses are located on campus at the Universidad Técnica de Machala in the city of Machala, Ecuador (3°15’ S, 79°57’ W), a region with abundant wild populations of *Ae. aegypti* and endemic arbovirus transmission. Experiments were conducted under ambient climate conditions (temperature range: 23.1—35.6° C, mean temperature: 28.4° C, relative humidity range: 43.9—95.0%, mean relative humidity: 75.8%). Each trial replicate was conducted with one control and one experimental house; the specific house used as the experimental or control house was alternated upon each replicate.

### Biological material

*Aedes aegypti* Linnaeus, 1762 eggs were provided by the Center for Research on Health in Latin America (CISeAL, by its Spanish acronym) at the Pontificia Universidad Católica del Ecuador. All strains used in this work originated from Ecuador, and had been maintained in laboratory conditions since 2015. Laboratory work was performed with strains originally collected in Ecuador from the cities of Guayaquil and Puerto Francisco de Orellana. Semi-filed work was performed with a strain originally collected in the city of Machala.

#### Semi-field experiments

Hatching and rearing of *Ae. aegypti* for semi-field experiments were performed at the Laboratory of Entomology at the Universidad Técnica de Machala. Considering this laboratory is located in a region where *Ae. aegypti* actively reproduces and thrives, environmental conditions (temperature: 28-32 °C; relative humidity: 60-80%) were not artificially controlled in our mosquito-rearing facilities. A vacuum pressure system was used to synchronize egg hatching (one hour exposure to obtain first stage larvae). Larvae were fed with finely ground fish food. At the pupal stage, males and females were separated. Adults were kept in 20×20×20 cm cages. Adults were fed on 10% sugar solution *ad libitum*. Each experimental semi-field experiment series used nulliparous females aged 1-5 days and starved for 24 hours prior to experimental release.

### Dried Attractive Bait Stations (DABS)

The DABS device consists of two concentric foam disks (an inner white disk one centimeter in diameter, and an outer black disk eight centimeters in diameter). Experimental DABS were impregnated with a 10% sucrose solution containing 1% boric acid as a lethal agent. Control DABS were impregnated with 10% sucrose solution without boric acid. Photographs of the devices are available in an additional file (see Supplementary Figure 2, Additional file 2), US Patent App. 15/990,931, 2018.

### Laboratory experiments

#### Series 1.1: Survival assessment of mosquitoes exposed to the device

To determine whether exposure to the DABS devices has an influence on adult mosquito survival probability, we conducted an experiment in which groups of 30 adult female mosquitoes, placed in a 15×15×15cm cage, were exposed during 48 hours to either a DABS device or a control device (sugar solution but no boric acid). We replicated each experiment four times. The assessment was repeated using each of the two laboratory strains described previously.

#### Series 1.2: Appraisal of biological mode of action of the device

To establish whether the toxic component of DABS needs to be ingested by the mosquitoes in order to exert its effect, we presented the devices to cohorts of adult females aged 1 to 7 days, which were unable to ingest food due to the surgical ablation of their mouthparts. To establish these cohorts, individual mosquitoes were first anesthetized by placing them at 4°C during 10-15 minutes. Anesthetized specimens were individually placed under a dissection microscope and, using a human hair, we tied a knot at the proboscis’ proximal end in order to create a constriction that would impede the flow of food. Subsequently, the part of the proboscis anterior to the knot was removed using micro-dissection scissor (Supplemental Figure 3). Following the surgery, mosquitoes were left to rest for 24 hours before being used in any experiment. To control for the potential negative effect of the anesthetizing procedure in mosquito survival, non-ablated mosquitoes used in control groups were also placed at 4°C during 10-15 minutes, and allowed to recover during 24 hours before experimental set-up.

We conducted the experiment with four separate cages, each with 20 starved mosquitoes (Table 2 shows different treatments). We treated cage 1 with toxic DABS devices and used 20 ablated mosquitoes; cage 2 held non-toxic control devices and 20 ablated mosquitoes. We treated cage 3 with toxic DABS devices and non-ablated mosquitoes; cage 4 held a non-toxic control device and non-ablated mosquitoes. We assessed mortality in all groups at 24 and 48 hours of exposure to the devices. We replicated the experiment three times.

We then conducted an experiment wherein 30 adult starved female mosquitoes aged 1 to 7 days were introduced to a cage with a DABS device, and 30 adult starved female mosquitoes of similar age were introduced to a cage with a non-toxic control device. We monitored cages for 24 hours, and removed dead mosquitoes by aspiration every hour from the cages. Using a dissection microscope, we removed the legs, head, and wings of every dead specimen onto a drop of 70% ethanol. Through this process we gently disrupted the abdominal cuticle to permit the exposure of internal tissues to the fixative. Afterwards we fixed individual mosquitoes in a solution containing 2.5% glutaraldehyde, 2.5% paraformaldehyde in 0.1M cacodylate buffer (pH 7.4), and stored them at 4°C for 72 hours. We then washed specimens in cacodylate buffer with 0.1M sucrose overnight. Post-fixing was achieved by leaving the specimens for two hours at 4°C in 2% osmium tetroxide in 0.1 cacodylate buffer (pH 7.4). Subsequently, individuals were stained using 2% uranyl acetate and left to rest for three hours in the dark at room temperature. Tissues were later dehydrated through a series of ethanol baths (50%, 70%, 95%, 100%). Afterwards, they were placed in propylene oxide for 30 minutes, then in a 1:1 volume propylene oxide resin mixture (Epon 812, Araldite 502, dodecenyl succinic anhydride, benzyl dimethylamine) for one hour and later, one more volume of resin was added and left on a rotator overnight. Finally, mosquitoes were embedded in resin and incubated at 60°C for 24 hours. Resin were stained using 2% uranyl acetate. We then utilized a transmission electron microscope to observe specimens and obtain micrographs of interesting tissues.

#### Series 1.3: Effects of the physiological status of the mosquitoes on the performance of DABS

We examined two different physiological statuses using mated starved female adult mosquitoes aged 1 to 7 days, namely blood fed and parous. We established females deemed as “blood fed” by selecting blood-engorged individuals immediately after a blood meal. We established females deemed as “parous” by first blood feeding and subsequently maintaining mosquitoes for 7 days under insectary conditions in order to ensure that they had oviposited before being used for experimentation. We set up two cages for each of the defined physiological statuses, each with 30 mosquitoes. One cage exposed the mosquitoes to an ATSB device, and the other held a control non-toxic device. We gathered survival data at 24 and 48 hours following introduction to the cages, and replicated these experiments three times.

#### Series 1.4: Shelf-life of the device

In order to determine the shelf life of ATSB devices, toxicity tests were performed using devices which had been stored for 38, 80, and 118 days after their production. For storage, devices were individually wrapped inside a sealed plastic bag and placed in an incubator at 28 ± 2°C and 80 ± 10% relative humidity. We conducted three replicates of previously described experiments for each storage time.

### Semi-field trials

#### Series 2.1: 24 hours of DABS exposure in experimental houses

Each house contained four DABS devices (control or treatment DABS as appropriate) suspended on strings attached to the roof of the house at a height of 30—50cm above ground and approximately 30cm from the nearest wall. Diagrams of the experimental layout are available in an additional file (see Supplementary Figure 3, Additional file 2). For each trial replicate, 50 female *Ae. aegypti* were released into each house through the escape window sleeve (release time 11:00a.m.— 2:00p.m.). Twenty-four hours after release, dead mosquitos were collected from the floor and window escape traps in each house, and the remaining live mosquitoes were captured with a hand-held aspirator (Prokopack, John W. Hock Company, Gainesville, USA). All live mosquitoes were labeled by experimental group and observed for 48 additional hours in laboratory cages (under laboratory conditions with food available). Mortality was calculated for 24 hours, 48 hours, and 72 hours. Six trial replicates were performed for Series 1.

#### Series 2.2: 48 hours of DABS exposure in experimental houses

Each house contained four DABS devices (control or treatment DABS as appropriate) and two sources of water (wet cotton in a black plastic bucket). Photographs of the water source are available in an additional file (see Supplementary Figure 3, Additional file 2). For each trial replicate, 50 female *Ae. aegypti* were released into each house through the escape window sleeve (release time 8:00a.m.—11:00a.m.). Forty-eight hours after release, dead mosquitos were collected in each house and remaining live mosquitoes were captured with an aspirator. Mortality was calculated for 48 hours. Six replicates were performed for Series 2.

#### Series 2.3: 48 hours of DABS exposure in experimental houses with competing attractant

Each house contained four DABS devices (control or treatment DABS as appropriate), two sources of water (wet cotton in a black plastic bucket), and 100 grams of peeled, cut apples in a dish placed on a chair in the center of the house as a competing attractant. Photographs of the competing attractant are available in an additional file (see Supplementary Figure 3, Additional file 2). Recently emerged female *Ae. aegypti* rely on sugar meals for energy, these meals may include aging fruit and female *Ae. aegypti* will feed on fructose (as is found in apples). For each trial replicate, 50 female *Ae. aegypti* were released into each house through the escape window sleeve (release time 9:00a.m.— 12:00p.m.). Forty-eight hours after release, dead mosquitos were collected in each house and remaining live mosquitoes were captured with an aspirator. Mortality was calculated for 48 hours. Six replicates were performed for Series 3.

### Statistical analyses

For the Series 1 experiments, data was processed, plotted, and analyzed using Python v2.7.13. For data processing we used Pandas v0.22.0 module. Plots were generated using the Plotly v3.10.0 module. We examined the normal distribution of the data with Kolmogorov-Smirnov and Shapiro-Wilk tests. In experiments in Series 1.1, 1.3, and 1.4 Student’s T-test comparisons were performed using the Scipy v1.0.0 module. In Series 1.2, one-way ANOVA was performed using Scipy v1.0.0 module with four experimental groups. Tukey’s range test, using Statsmodels v.0.10.0 module, was performed after ANOVA for determining ranges for each group. All data and codes used for this data have been stored in a private online git repository and will be provided upon request. In Series 2.1— 2.3, mosquito mortality data from each series were compared using a two-tailed paired t-test (paired by replicate). Mean mosquito mortality was compared across series using a two-tailed t-test. Data were analyzed using Excel (Microsoft, Redmond, USA).

## List of abbreviations

*Ae*. aegypti: *Aedes aegypti* Linnaeus, 1762
*Ae. albopictus*: *Aedes albopictus* Skuse 1894
ATSB: attractive toxic sugar bait
DABS: dried attractive bait stations
cm: centimeters
m: meters

## Declarations

### Ethics approval

These study protocols were found to be exempt from Institutional Review Board review by Syracuse University Institutional Review Board.

### Consent for publication

Not applicable

### Availability of data and material

The datasets used and/or analysed during the current study are available from the corresponding author on reasonable request.

### Author contributions

MN, ASI, and DL designed the study.

MN, ASI, DL, and EBA provided study materials.

GR, ML, AA, BM VS and FHH performed the experiments.

RH performed electron microscopy work

RS and GR conducted data analysis.

RS, GR, MN, VS, and FHH drafted the manuscript.

All authors reviewed and approved the final draft of the manuscript.

## Acknowledgements

The authors would like to thank Cinthya Cueva Aponte for her assistance with study documentation, Chad Ward and Matt Kovacs for assistance with mosquito rearing, Mario Grijalva for his logistical support for the performance of electron microscopy, Felipe Andrade and Phillip Huegli for providing useful recommendations on programming, and Santiago Cadena for his support in insectary maintenance.

## Additional files

Additional file 1 (pdf)

Supplementary Figure 1: Experimental Houses

Additional file 2 (pdf)

Supplementary Figure 2: Experimental Layout

Additional file 3 (pdf)

Supplemental Figure 2: Feeding disruption procedure. A) Anesthetized individual with whole proboscis. B) Human hair tied at the proximal end of the proboscis. C) Micro-dissection scissor removal of proboscis’ segment anterior to the knot. D) Feeding disrupted individual.

## Supplemental table

## References

1. PAHO, WHO. Epidemiological Update: Dengue. 2019;1–14.

2. Boyer S, Calvez E, Chouin-Carneiro T, Diallo D, Failloux AB. An overview of mosquito vectors of Zika virus. Microbes Infect. 2018;20:646–60.

3. Olliaro P, Fouque F, Kroeger A, Bowman L, Velayudhan R, Santelli AC, et al. Improved tools and strategies for the prevention and control of arboviral diseases: A research-to-policy forum. PLoS Negl Trop Dis. 2018;12:e0005967.

4. Oxborough RM. Trends in US President’s Malaria Initiative-funded indoor residual spray coverage and insecticide choice in sub-Saharan Africa (2008-2015): urgent need for affordable, long-lasting insecticides. Malar J. 2016;15:146.

5. Harrington LC, Scott TW, Lerdthusnee K, Coleman RC, Costero A, Clark GG, et al. Dispersal of the dengue vector Aedes aegypti within and between rural communities. Am J Trop Med Hyg. 2005;72:209–20.

6. Hemingway J, Ranson H. Insecticide Resistance in Insect Vectors of Human Disease. Annu Rev Entomol. 2000;45:371–91.

7. Chanda E, Kamuliwo M, Sikaala CH, Thomsen EK, Ranson H, Hemingway J, et al. An operational framework for insecticide resistance management planning. Emerg Infect Dis. 2016;22:773–9.

8. Ryan SJ, Mundis SJ, Aguirre A, Lippi CA, Beltrán E, Heras F, et al. Seasonal and geographic variation in insecticide resistance in Aedes aegypti in southern Ecuador. PLoS Negl Trop Dis. 2019;13:e0007448.

9. Kraemer MUG, Sinka ME, Duda KA, Mylne A, Shearer FM, Barker CM, et al. The global distribution of the arbovirus vectors Aedes aegypti and Ae. albopictus. Elife. 2015;4:e08347.

10. Yuval B. The other habit: sugar feeding by mosquitoes. Bull Soc Vector Ecol. Society for Vector Ecology; 1992;17:150–6.

11. Foster WA. Mosquito sugar feeding and reproductive energetics. Annu Rev Entomol. 1995;40:443–74.

12. Fiorenzano JM, Koehler PG, Xue R De. Attractive toxic sugar bait (ATSB) for control of mosquitoes and its impact on non-target organisms: A review. Int J Environ Res Public Health. 2017;14.

13. Tenywa FC, Kambagha A, Saddler A, Maia MF. The development of an ivermectin-based attractive toxic sugar bait (ATSB) to target Anopheles arabiensis. Malar J. 2017;16:1–10.

14. Maia MF, Tenywa FC, Nelson H, Kambagha A, Ashura A, Bakari I, et al. Attractive toxic sugar baits for controlling mosquitoes: A qualitative study in Bagamoyo, Tanzania. Malar J. 2018;17:1–6.

15. Stewart ZP, Oxborough RM, Tungu PK, Kirby MJ, Rowland MW, Irish SR. Indoor application of attractive toxic sugar bait (ATSB) in combination with mosquito nets for control of pyrethroid-resistant mosquitoes. PLoS One. 2013;8.

16. Qualls WA, Müller GC, Revay EE, Allan SA, Arheart KL, Beier JC, et al. Evaluation of attractive toxic sugar bait (ATSB)-Barrier for control of vector and nuisance mosquitoes and its effect on non-target organisms in sub-tropical environments in Florida. Acta Trop. 2014;131:104–10.

17. Qualls WA, Müller GC, Traore SF, Traore MM, Arheart KL, Doumbia S, et al. Indoor use of attractive toxic sugar bait (ATSB) to effectively control malaria vectors in Mali, West Africa. Malar J. Malaria Journal; 2015;14:301.

18. Qualls WA, Scott-Fiorenzano J, Müller GC, Arheart KL, Beier JC, Xue R-D. Evaluation and Adaptation of Attractive Toxic Sugar Baits For Culex tarsalis and Culex quinquefasciatus Control In The Coachella Valley, Southern California. J Am Mosq Control Assoc. 2017;32:292–9.

19. Müller GC, Junnila A, Schlein Y, Ller GCM. Effective Control of Adult Culex pipiens by Spraying an Attractive Toxic Sugar Bait Solution in the Vegetation Near Larval Habitats. Source J Med Entomol J Med Entomol. 2010;47:63–6.

20. Revay EE, Müller GC, Qualls WA, Kline DL, Naranjo DP, Arheart KL, et al. Control of Aedes albopictus with attractive toxic sugar baits (ATSB) and potential impact on non-target organisms in St. Augustine, Florida. Parasitol Res. 2014;113:73–9.

21. Junnila A, Revay EE, Müller GC, Kravchenko V, Qualls WA, Xue R de, et al. Efficacy of attractive toxic sugar baits (ATSB) against Aedes albopictus with garlic oil encapsulated in beta-cyclodextrin as the active ingredient. Acta Trop. 2015;152:195–200.

22. Scott-Fiorenzano JM, Fulcher AP, Seeger KE, Allan SA, Kline DL, Koehler PG, et al. Evaluations of dual attractant toxic sugar baits for surveillance and control of Aedes aegypti and Aedes albopictus in Florida. Parasites and Vectors. 2017;10:1–9.

23. Scott JM, Seeger KE, Gibson-Corrado J, Muller GC, Xue R De. Attractive Toxic Sugar Bait (ATSB) Mixed With Pyriproxyfen for Control of Larval Aedes albopictus (Diptera: Culicidae) Through Fecal Deposits of Adult Mosquitoes. J Med Entomol. 2017;54:236–8.

24. Khallaayoune K, Qualls WA, Revay EE, Allan SA, Arheart KL, Kravchenko VD, et al. Attractive Toxic Sugar Baits: Control of Mosquitoes With the Low-Risk Active Ingredient Dinotefuran and Potential Impacts on Nontarget Organisms in Morocco. Environ Entomol. 2013;42:1040–5.

25. Lea AO. Sugar-baited insecticide residues against mosquitoes. Mosq News. 1965;25:65.

26. Fikrig K, Johnson BJ, Fish D, Ritchie SA. Assessment of synthetic floral-based attractants and sugar baits to capture male and female Aedes aegypti (Diptera: Culicidae). Parasites and Vectors. 2017;10:1–9.

27. Xue R-D, Ali A, Kline DL, Barnard DR. Field Evaluation of Boric Acid- and Fipronil-Based Bait Stations Against Adult Mosquitoes. J Am Mosq Control Assoc. 2008;24:415–8.

28. Martinez-Ibarra JA, Rodriguez MH, Arredondo-Jiménez JI, Yuval B, Arredondo-Jimenez JI, Yuval B. Influence of plant abundance on nectar feeding by Aedes aegypti (Diptera: Culicidae) in southern Mexico. J Med Entomol. 1997;34:589–93.

29. Spencer CY, Pendergast TH, Harrington LC. Fructose variation in the dengue vector, Aedes aegypti, during high and low transmission seasons in the Mae Sot region of Thailand. J Am Mosq Control Assoc. 2005;21:177–81.

30. van Handel E, Edman JD, Day JF, Scott TW, Clark GG, Reiter P, et al. Plant-sugar, glycogen, and lipid assay of Aedes aegypti collected in urban Puerto Rico and rural Florida. J Am Mosq Control Assoc. 1994;10:149–53.

31. Sissoko F, Junnila A, Traore MM, Traore SF, Doumbia S, Dembele SM, et al. Frequent sugar feeding behavior by Aedes aegypti in Bamako, Mali makes them ideal candidates for control with attractive toxic sugar baits (ATSB). PLoS One. 2019;14:1–21.

32. Xue R-D, Kline DL, Ali A, Barnard DR. Application of boric acid baits to plant foliage for adult mosquito control. J Am Mosq Control Assoc. American Mosquito Control Association; 2006;22:497–500.

33. Xue R-D, Barnard DR. Boric acid bait kills adult mosquitoes (Diptera: Culicidae). J Econ Entomol. 2003;96:1559–62.

34. Garcia-Rejon J, Loroño-Pino MA, Farfan-Ale JA, Flores-Flores L, Del Pilar Rosado-Paredes E, Rivero-Cardenas N, et al. Dengue virus-infected Aedes aegypti in the home environment. Am J Trop Med Hyg. 2008;79:940–50.

35. Kuno G. Review of the factors modulating dengue transmission. Epidemiol Rev. 1995;17:321–35.

36. Fay RW, Prince WH. A modified visual trap for Aedes aegypti. Mosq News. 1970;

37. Barrera R, Mackay AJ, Amador MA. An improved trap to capture adult container-inhabiting mosquitoes. J Am Mosq Control Assoc. 2013;29:358–68.

38. Meunier N, Marion-Poll F, Rospars JP, Tanimura T. Peripheral coding of bitter taste in Drosophila. J Neurobiol. 2003;56:139–52.

39. Dennis EJ, Goldman O V., Vosshall LB. Aedes aegypti Mosquitoes Use Their Legs to Sense DEET on Contact. Curr Biol. 2019;29:1551-1556.e5.

40. Hemme RR, Thomas CL, Chadee DD, Severson DW. Influence of urban landscapes on population dynamics in a short-distance migrant mosquito: evidence for the dengue vector Aedes aegypti. PLoS Negl Trop Dis. 2010;4:e634.

41. Heydari N, Larsen D, Neira M, Beltrán Ayala E, Fernandez P, Adrian J, et al. Household Dengue Prevention Interventions, Expenditures, and Barriers to Aedes aegypti Control in Machala, Ecuador. Int J Environ Res Public Health. 2017;14:196.

